# Name Conditioning in Event-Related Brain Potentials

**DOI:** 10.1101/194134

**Authors:** Boris Kotchoubey, Yuri G. Pavlov

**Author notes:** Corresponding author: B. Kotchoubey, Institute of Medical psychology and Behavioral Neurobiology, University of Tübingen, Silcherstr. 5, 72076 Tübingen, Germany. Conflict of interests: None.

## Abstract

Four experiments are reported in which two harmonic tones (CS+ and CS-) were paired with a participant’s own name (SON) and different names (DN), respectively. A third tone was not paired with any other stimulus and served as a standard (frequent stimulus) in a three-stimuli oddball paradigm. The larger posterior positivity (P3) to SON than DN, found in previous studies, was replicated in all experiments. Conditioning of the P3 response was albeit observed in two similar experiments (1 and 3), but the obtained effects were weak and not identical in the two experiments. Only Experiment 4, where the number of CS/UCS pairings and the Stimulus-Onset Asynchrony between CS and UCS were increased, showed clear CS+/CS- differences both in time and time-frequency domains. Surprisingly, differential responses to CS+ and CS- were also obtained in Experiment 2, although SON and DN in that experiment were masked and never consciously recognized as meaning words (recognition rate 0/63 participants). The results are discussed in the context of other ERP conditioning experiments and, particularly, the studies of non-conscious effect on ERP. Several further experiments are suggested to replicate and extend the present findings and to remove the remaining methodological limitations.

---

Effects of classical conditioning on human Event-Related Brain Potentials (ERPs) have been examined in a number of studies (for review, see Christoffersen & Schachtman, 2016; Miskovic & Keil, 2012). Most of them used highly aversive unconditioned stimuli (UCS) (e.g., Hermann, Ziegler, Birbaumer, & Flor, 2000; Pizzagalli, Greishar, & Davidson, 2003). As a rule, CS are complex visual stimuli, e.g., faces (Begleiter, & Platz, 1969) or words (Montoya, et al., 1996). Very few studies employed both CS and UCS of auditory modality (Hugdahl & Norby, 1991; Heim & Keil, 2006; Pauli & Röder, 2008; Juan et al., 2016).

Because auditory classical conditioning, due to its technical simplicity, can be applied in children and severely disabled individuals, and because using highly aversive UCS in these groups is ethically problematic, looking for other kinds of UCS is important. Having in mind the potential application in patients with severe brain damage, in the present study we intended to explore the effect of classical conditioning on ERP to simple stimuli. Relatively simple harmonic tones were chosen as CS, based on the finding that harmonic tones elicit more distinct and stable ERP effects than sine tones in both healthy individuals (Tervaniemi et al., 2000) and neurological patients (Kotchoubey et al., 2003). An individual’s own name, which has been suggested to possess particular significance for the individual, was used as a non-aversive UCS. The effects of a subject’s own name (SON) on ERPs have been established in normal populations (Fischler et al., 1987) and severely brain-injured patients (Perrin et al., 2006), in waking state (Holeckova et al., 2006) and during sleep (Perrin, Garcia-Larrea, Mauguiere, & Bastuji, 1999). In a three-stimulus oddball, in which SON and a control stimulus (usually, a different name: DN) are presented as two rare stimuli, SON elicits a larger P3 component than DN (e.g., Perrin et al., 2006; Kotchoubey et al., 2004). We expected to obtain a similar effect in response to harmonic tones (presented as CS) paired with names.

## Methods: General

Three different groups of healthy participants took part in the study: one group (nine males and 14 females, aged 22-29) in Experiment 1, the second group (nine males and 13 females, aged 22-29) in Experiments 2 and 3, and the third group (twelve males and 13 females, aged 19-42) in Experiment 4. In the second group Experiment 2 always preceded Experiment 3. Data of two males in Experiment 2 were excluded (thus the group contained 7 males).

None of the participants had had any disease of the nervous system or hearing disorders in the past, or reported use of any drugs during the last week before the experiment. Participants were seated in a comfortable chair and asked to close their eyes and to listen attentively to the stimuli. Informed consent was obtained from each participant. The study was approved by the Ethical Committee of the University of Tübingen.

The EEG in all experiments was recorded using 64 active ActiCHamp electrodes (Easycap GmbH, Herrsching, Germany) located according to the extended 10-20 system. The vertical and horizontal electrooculagram were also recorded. The resistance was below 15 kOhm. Online reference was at Cz, offline re-referenced to average mastoids. The digitalization rate was 1000 Hz.

Off-line inspection of the recordings revealed in some traces poor data quality in one or two of the 64 channels. These channels were replaced with interpolation of the adjacent electrodes. After this, an Independent Component Analysis (ICA) was employed for each participant to separate and remove activity due to ocular artifacts using the AMICA algorithm (Palmer et al., 2012). Components clearly related to eye movements were removed using EEGLAB. Additionally, components that were mapped onto one electrode and could be clearly distinguished from EEG signals were subtracted from the data. EEG segments that still contained artifacts after ICA correction were dismissed. The ERPs were filtered within a band from 0.1 to 30 Hz and averaged in relation to a baseline from -200 ms to 0 ms. As we supposed that the responses would change during the roughly 7-minute test phase, ERPs were averaged separately for the first, second, and third thirds of the whole sequence of 400 stimuli. The three periods, corresponding to stimuli 1 – 133, stimuli 134 – 266, and stimuli 267 – 400, will be referred to as T1, T2, and T3, respectively. Each average included at least 18 (usually 20) CS+ and CS-.

The amplitudes of ERP components were measured as the area under the curve within the time windows slightly different for UCS (N1: 70-120 ms, P3a: 240-300 ms, P3b: 300-400 ms, Late Time Window [LTW]: 400-600 ms) and CS (N1: 100-140 ms, P3a: 220-280 ms; P3b: 280-380 ms; LTW: 380-550 ms). The LTW was not designated as “P” or “N” because the amplitude was negative in anterior but positive in posterior leads. The time-frequency analysis was performed using Morlet wavelet by means of the Fieldtrip toolbox and followed the method of Cavanagh et al. (2010). The entire epochs were defined as [-1500 2500] ms to avoid edge artifacts. Baseline correction and decibel normalization was performed in respect to [-400 -100] ms interval. After a visual exploration of grand averages across all subjects, conditions, channels, and experiments, two time-frequency (TF) windows were extracted: “P3” (200-350 ms, 9-12 Hz) and “LTW” (400-650 ms, 6-8 Hz).

For brevity, the present report describes only those data that are related to the critical comparison between the CS+ and CS- responses at the midline electrodes Fz, Cz, and Pz, were the effects were best pronounced. The statistical analysis was performed using a repeated measures ANOVA with factors Stimulus, Site, and Time. When appropriate, we used Greenhouse-Geisser non-sphericity correction for degrees of freedom.

## Experiment 1

### Methods

ERPs were recorded to three chords, each consisting of five harmonic tonal frequencies (e.g., 330, 660, 1320, 2640, and 5280 Hz). One of the chords was used as standard and the other two served as CS+ and CS-. During the *acquisition phase*, CS+ was paired 21 times with SON, and CS- was randomly paired 21 times with three different names (DN). All names were spoken with the official German pronunciation by a female speaker, not familiar to any participant. The control names originated from the same pool of the most frequent German names used for each subject’s own name. They always had a very similar duration as the own name (means 669 ms and 676 ms) and contained the same number of syllables. The standard was presented 21 times, not accompanied by any other stimulus. Tone duration was 200 ms, and the intensity was 75 dB above the average threshold. The stimulus-onset-asynchrony (SOA) within a pair tone-name was 300 ms. The SOA after a tone-word pair was 1700-1800 ms, and after standards it was 1150-1250 ms. All stimuli were presented binaurally through aerodynamic earphones, in a pseudorandomized order, in which none of the three tones appeared more than three times in a row.

In the *test phase*, which followed immediately after the *acquisition phase*, the standard was presented 280 times, and CS+ and CS- 60 times each. No other stimuli were presented. The SOA varied between 950 and 1050 ms. The order of presentation was randomized except that CS+ and CS- could not be delivered more than twice in a row.

### Results

All participants reported after the experiment, that they had heard “two or three” different harmonic tones, and that at the beginning one of the tones was linked to their own name, and another tone, to other names.

The *acquisition phase* replicated the already known effect of a larger P3 to SON. The effect was most clear in the P3a window (F(1,22) = 11.55, p = .003, η^2^ = .34). Also in the LTW, the amplitude was negative to DN but positive to SON at Cz and Pz, yielding a significant Stimulus x Site interaction: F(2,44) = 5.37, p = .014, η^2^ = .20. Importantly, the P3a amplitude was larger to CS+ than CS- (mean amplitudes 3.47 versus 1.78 µV; F(1,22) = 10.63, p = .004, η^2^ = .33). The P3(a) effect, was, however, instable and disappeared in the *test phase*, in which no differences between CS+ and CS- responses were observed. The average ERP waveforms are shown in Figures 1 and 2.

**Figure 1.**
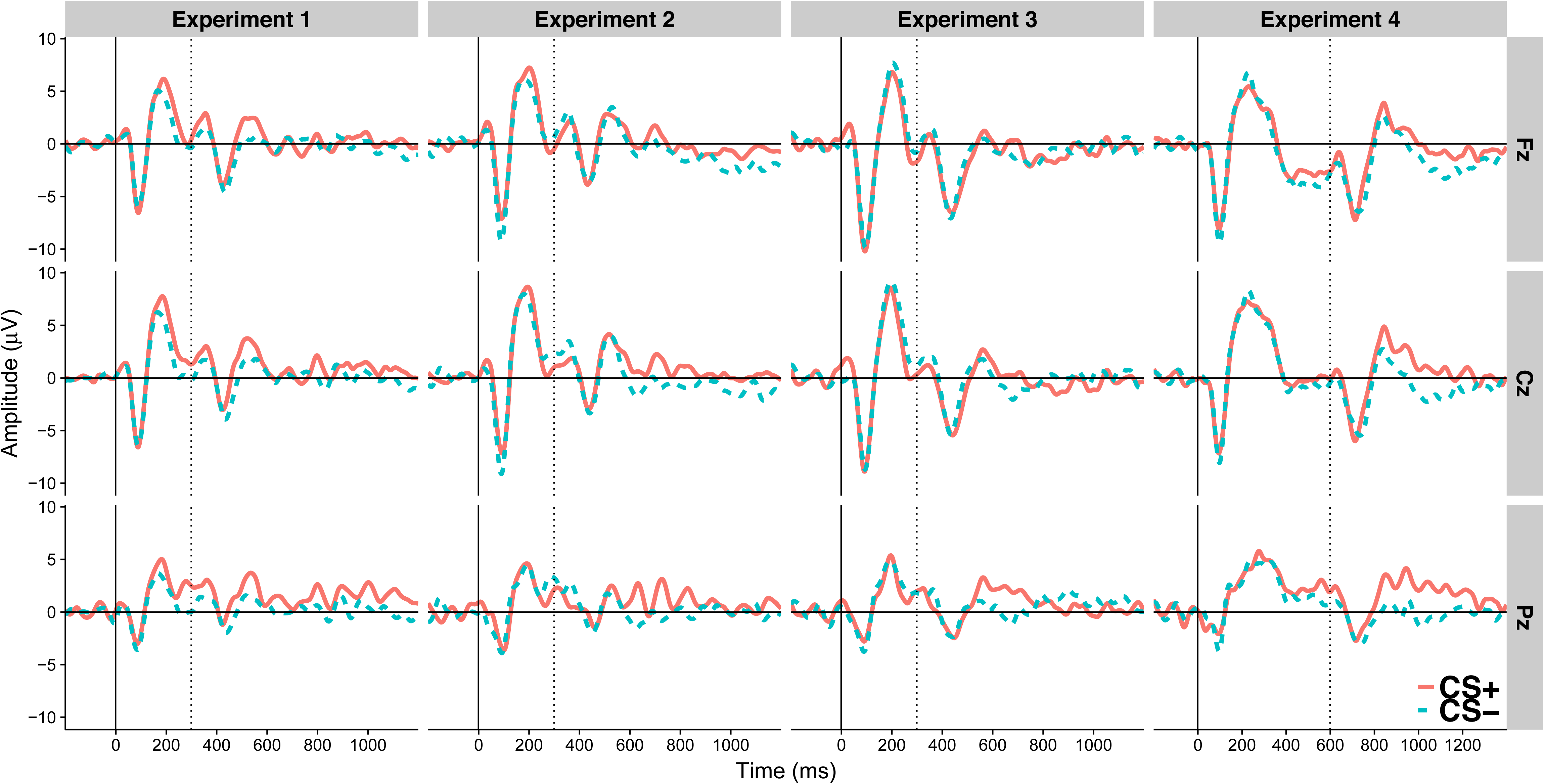
Grand average ERP obtained in the acquisition phase of the four experiments in response to CS+/SON (solid line) and CS-/DN (dashed line) combinations. Solid vertical lines indicate the onset of CS, and dotted vertical lines, the onset of UCS. Note that the rightmost column shows ERP averaged across the first one-third of Experiment 4, to make the number of trials comparable for all experiments. Positivity is plotted upwards.

**Figure 2.**
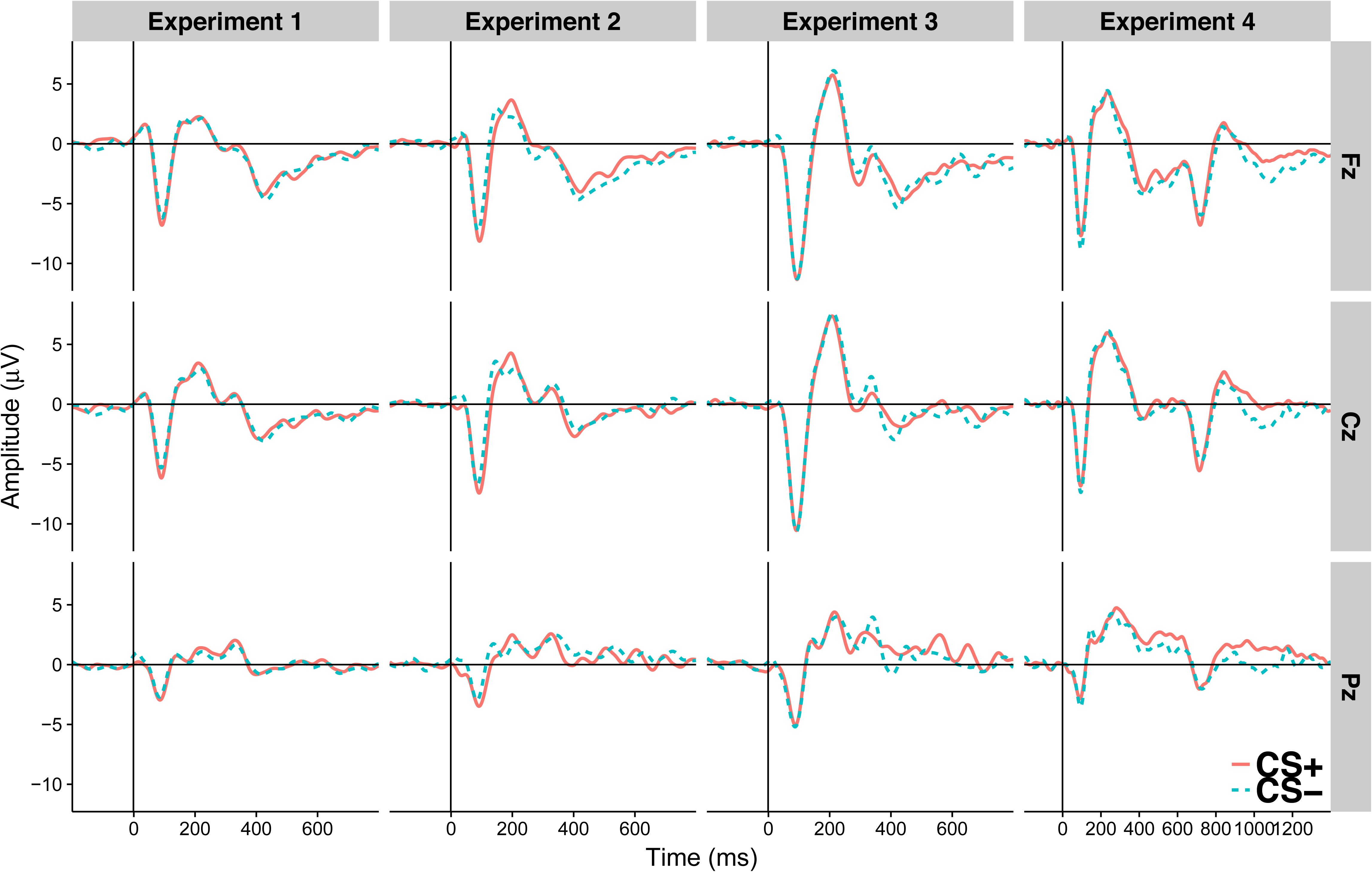
Experiment 1 to 3: Grand average ERP waveforms obtained in the test phase. Experiment 4: Grand average across the whole experiment. The number of included trials is similar for all four experiments. Labelings are similar to Figure 1.

## Experiment 2

### Methods

In Experiment 2 the names were completely masked while preserving their acoustical features. The first 25% of time points of an original name were multiplied by a linearly spaced vector of coefficients from 1.5 to 0, and the remaining 75% points were set to 0. Then, the first 25% of time points of the same name played backwards were multiplied by a linearly spaced vector of coefficients from 0 to 1.5, and the last 75% time points remained unchanged. Finally, the two files were added. This technique permitted to attain the same intensity-by-time dynamics as in the original names. In a pilot experiment the stimuli were presented to forty healthy participants. None of them was able to recognize any name including their own.

### Results

Participants reported that they had heard “two or three” different harmonic tones, and that at the beginning of the stimulation also other stimuli had been presented that sounded like non-comprehendible words of an exotic (non-European) language.

Despite the lack of subjectively perceived differences between the two UCS, P3b had a larger amplitude to SON than DN, particularly at Pz (main Stimulus effect: F(1,20) = 5.04, p = .035, η^2^ = .20; Stimulus x Site interaction: F(2,40) = 5.85, p = .017, η^2^ = .23). P3a, in contrast to Experiment 1, did not differ between SON and DN. During the *test phase*, N1 and P3a were significantly larger to CS+ than CS- (F(1,20) = 20.21, p < .001, η^2^ = .50; and 4.95, p = .038, η^2^ = .20, for N1 and P3a, resp.).

## Experiment 3

### Methods

Although CS+ and CS- differed in the perceived pitch, in Experiment 3 they also were delivered monaurally in two different ears to further increase their discriminability. The side of presentation was counterbalanced among the participants.

### Results

Participants reported after the experiment that they had heard three different harmonic tones. They were aware of the association between tones and names.

Both P3a and P3b were larger to SON than DN at Pz (Stimulus x Site interaction: F(2,42) = 4.54, p = .03, η^2^ = .18; and 4.75, p = .026, η^2^ = .18; for P3a and P3b, resp.). The main effect of Stimulus at Pz was significant (p < .02) for both components. The general main Stimulus effect approached significance for P3b: F(1,21) = 3.87, p = .058, η^2^ = .15). In the *test phase*, the amplitude in the LTW was more positive to CS+ than CS- (F(1,21) = 4.68, p =.042, η^2^ = .18). P3a, in contrast to Experiments 1 and 2, appeared to be slightly larger to CS- than CS+, but this difference did not attain significance.

## Experiment 4

### Methods

This Experiment entailed only one phase, during which the standard was presented 280 times, and CS+ and CS- 60 times each. Tone duration was 100 ms including 5 ms rise/fall phase. All CS were followed by the corresponding UCS with SOA of 600 ms. This design aimed at the recording of the late ERP components to CS immediately during acquisition, because in the other experiments (with SOA of 300 ms) they could be measured only in the test phase.

### Results

SON elicited a larger P3b than DN (F(1,24) = 5.08, p = .034, η^2^ = .18), and this difference steadily decreased with time (Stimulus x Time interaction: F(2,48) = 3.40, p = .040, η^2^ = .13). The LTW was consistently negative to DN, but negative at Fz and positive in T1 at Pz (thus zero on average) to SON, resulting in a main effect of Stimulus (F(1,24) = 8.68, p = .007, η^2^ = .27) and a Stimulus x Site x Time interaction (F(4,96) = 3.51, p = .023, η^2^ = .13).

CS+ elicited more positive P3b (F(1,24) = 4.40, p = .047, η^2^ = .16) and more positive LTW amplitude (F(1,24) = 7.83, p = .01, η^2^ = .25) than CS-. When only the first 20 CS+/SON and CS-/DN pairs are taken into the analysis (for comparison with the other experiments in which CS+/SON and CS-/pairs were presented 20 times each), the same difference is also significant for P3a: F(1,24) = 5.36, p = .03, η^2^ = .18.

### Time-frequency analysis

As can be seen in Figure 3, the upper theta activity in the LTW in response to the tones was suppressed as compared with the baseline. In experiments 2 and 4, this suppression was stronger to CS- than CS + at Pz (main effect of Stimulus in Experiment 2: F(1,20) = 6.12, p = .02, η^2^ = .23; Stimulus x Site interaction in Experiment 4: F(2,42) = 3.52, p = .05, η^2^ = .13; main effect of Stimulus at Pz: F(1,24) = 4.69, p = .03, η^2^ = .16). Two further effects were observed in Experiment 4. First, alpha activity in the P3 window was desynchronized after CS+ but not after CS- (main effect of Stimulus: F(1,24) = 4.67, p = .04, η^2^ = .16). Second, theta activity was strongly suppressed in the LTW at Pz after DN, but less after SON (Stimulus x Site interaction: F(2,42) = 3.76, p = .04, η^2^ = .14).

**Figure 3.**
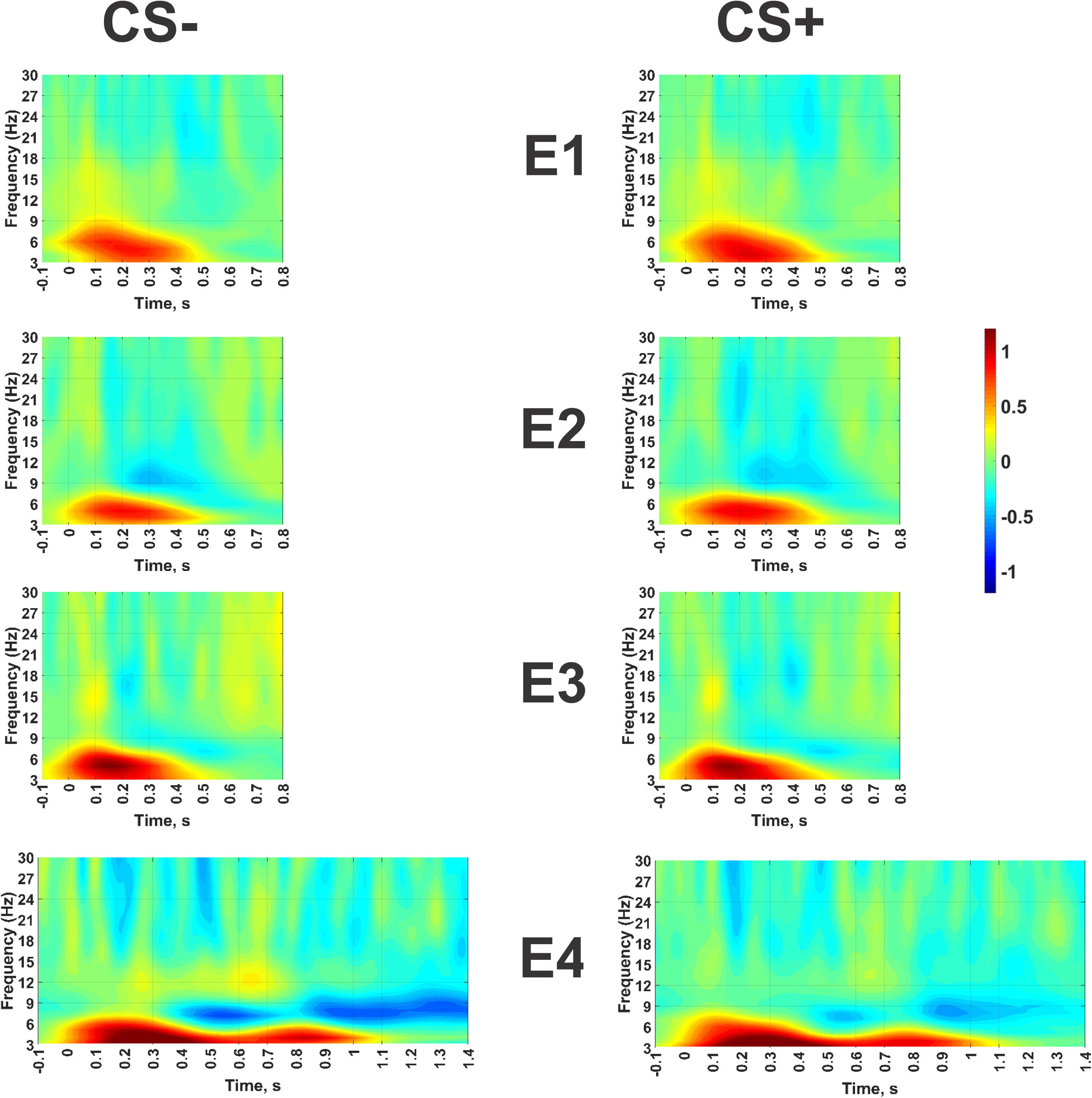
Results of the time-frequency analysis. Blue colors indicate power decrease, and yellow/red colors indicate power increase, as compared with the baseline. The color scale is presented in dB (that is, a value of +10 dB would mean the increase of the EEG power in ten times). Note that the epochs are broader than those shown in Figures 1 and 2, because the time-frequency analysis has, by definition, a lower temporal resolution than the analysis in time domain. No significant effects were found in Experiments 1 and 3.

## Discussion

The results are an intriguing mix of expected and surprising findings. First, we replicated the already known fact of an increased P3 to SON as compared with DN in a different paradigm. The increase was observed in all three experiments in which participants were able to recognize their own names. We hypothesized that the effect would be partially transferred to the related CS. In fact, CS+ elicited a larger positivity in the P3a time window than CS- during acquisition in Experiments 1 and 4, but not in Experiment 3 where it was expected as well. We also expected that the stable reinforcement in Experiment 4 would result in an increase of the difference between CS+ and CS-, and this hypothesis was confirmed. Due to the short CS/UCS SOA we could not record ERP components later than P3a in the acquisition phase of Experiments 1-3. In Experiment 4, with a longer SOA, we found that the short-living effect in the P3a time window stabilized and extended to the later time intervals between 300 and 600 ms.

To summarize, these findings indicate that classical conditioning of electrophysiological responses using the own name as UCS is possible. The differential responses to CS+ and CS- in Experiment 4 were demonstrated in both time (i.e., late ERP positivities) and time-frequency domains (i.e., higher theta but lower alpha activity to CS+ than CS-). Importantly, the better differentiation between the two CS was found in the same experiment in which the two UCS were also best distinguished. Generally, however, learning effects were relatively weak and unstable. Experiment 4, in which the clearest learning effects were obtained, did not contain extinction. The only effect that survived extinction was the differential response in the LTW in Experiment 3. Therefore, we do not believe that the paradigm can be used in clinical applications in its present form; rather, further methodological work is needed. Another limitation concerns the theoretical interpretation. Although the effect was primarily concentrated around P3a, we cannot rule out that other ERP components (e.g., P2 or P3b) also contributed to its generation. Also this issue should be followed in further experiments.

On the background of these largely predicted findings, the results of Experiment 2 were fully unexpected. Remember the masked names from this experiment were presented to a total of 63 individuals (40 participants of the pilot experiment + 22 participants of Experiment 2 + the first author who was unaware about masking developed by the second author), and none of them (0/63) recognized any name including their own. The stimuli were not even perceived as meaningful words. ERP, however, demonstrated a significantly larger positivity to SON than DN. Notably, this differential response was about 100 ms later than in all three experiments where SON was consciously recognized, indicating the presence of some additional processing operation(s). Although conditional responses in Experiment 2 did not differ during acquisition, the test phase revealed a significant P3a effect, quite similar to that in the acquisition phase of Experiment 1 and 4, and additionally, a very strong (η^2^ = .50) enhancement of the N1 amplitude to CS+ as compared with CS-. This N1 effect was very consistent at the individual level, being observed in 20 of the 22 participants.

During acquisition, SON was presented as UCS+ with the same frequency as three DN as UCS-. Therefore, each individual DN was presented three times less frequently than SON. Although this fact is a methodological limitation of the present study, it can hardly be responsible for any expected or surprising result. Firstly, this arrangement was the same in all four experiments, but the results were different. Secondly, N1 and P3a are expected to be larger to rare than frequent stimuli. If N1 is superposed by a Mismatch Negativity, this wave is also larger to rare than frequent stimuli. On this basis we might expect larger amplitudes to CS- (previously linked to rare DN) than to CS+ (previously linked to the more frequent SON), but the opposite was found. Only the *non-significant* increase of P3a in Experiment 3 would be in line with the frequentist interpretation. But even in this case such interpretation meets a considerable problem: it is fully unclear why ERP responses differ between the two CS having *equal frequencies* (and only linked to stimuli of different frequencies), whereas these responses *do not differ* between the two UCS having *different* frequencies.

If explanations related to experimental methodology are rejected, the only possible interpretation of the data of Experiment 2 remains that participants subconsciously distinguished between SON and DN even though they did not recognize them. This is particularly possible because our masking technique assured the similar intensity/time function of unmasked and masked UCS. If, for example, a sound file consisted of 100 data points, the first point was the same in masked and unmasked stimuli, the second point contained 95.8% common information, the third point 91.7%, etc. It may, therefore, be speculated that participants unconsciously recognized different personal significance of the stimuli even though they did not identify their content.

The fact that stimuli that are not consciously recognized can nonetheless elicit significant ERP effects has been shown in numerous studies (reviews Shevrin, 2001; Dehaene et al., 2006). A discussion of whether *all* kinds of learning in the brain can happen outside awareness (e.g., Hassin, 2013) will go far beyond the topic of classical conditioning, where subliminal effects on ERPs have repeatedly been shown, most recently by Beckes, Coan and Morris (2013). Most such subliminal effects are similar to (but usually weaker than) the corresponding effect of supraliminal, consciously perceived stimuli, while in the current study the results of Experiment 2 differ qualitatively from those of other experiments. A direct comparison between different studies is hardly possible due to the huge variability of techniques making stimuli non-recognizable. However, some studies demonstrated non-conscious ERP effects different from, stronger or even faster than similar effect to consciously perceived stimuli (e.g., Dehaene et al., 1998; Eimer & Schlaghecken, 1998; Williams et al., 2004). For instance, Eimer and Schlaghecken (1998) showed nearly equally strong, but opposite in direction, effects of masked and unmasked stimuli on the Lateralized Readiness Potential, an ERP index of motor preparation. All these and similar findings pose serious theoretical problems for both strong and weak single-process models of classical conditioning discussed by Lovibond and Shanks (2002) and favor their third, dual-process model.

Taking into account that this is the first indication of conditioning effects of a subject’s own name, suggestions for further experiments can be made. First, an extensive acquisition phase like in our Experiment 4 should be combined with a test phase like in Experiments 1 and 3. Second, an additional experiment using one DN should clarify the issue of stimulus frequencies. Third, regarding the putative non-conscious conditioning, different kinds of masking should be compared in respect of their ERP effects, and sophisticated techniques of debriefing should be used to explore the possible role of poorly formulated affective features of stimuli.

## Acknowledgement

The study was supported by the German Research Society (Deutsche Forschungsgemeinschaft), Grant KO-1753/13. We thank Ms. Janna Holst for her help in running the experiments.

